# Genetic Variability of Human Angiotensin-Converting Enzyme 2 (hACE2) Among Various Ethnic Populations

**DOI:** 10.1101/2020.04.14.041434

**Authors:** Quan Li, Zanxia Cao, Proton Rahman

**Affiliations:** Department of Medicine, Faculty of Medicine, Memorial University, St. John’s, Newfoundland, Newfoundland and Labrador, Canada A1B 3X9; Princess Margaret Cancer Centre, University Health Network, University of Toronto, Toronto, Ontario, Canada M5S 1A8; Shandong Provincial Key Laboratory of Biophysics, Institute of Biophysics, Dezhou University, Dezhou 253023, China

**Keywords:** Covid-19, SARS-CoV-2, Angiotensin-converting enzyme 2 (ACE2), gene expression, ethnic variation

## Abstract

There appears to be large regional variations for susceptibility, severity and mortality for Covid-19 infections. We set out to examine genetic differences in the human angiotensin-converting enzyme 2 (hACE2) gene, as its receptor serves as a cellular entry for SARS- CoV-2. By comparing 56,885 Non-Finnish European and 9,197 East Asians (including 1,909 Koreans) four missense mutations were noted in the hACE2 gene. Molecular dynamic demonstrated that two of these variants (K26R and I468V) may affect binding characteristics between S protein of the virus and hACE2 receptor. We also examined hACE2 gene expression in eight global populations from the HapMap3 and noted marginal differences in expression for some populations as compared to the Chinese population. However, for both of our studies, the magnitude of the difference was small and the significance is not clear in the absence of further in vitro and functional studies.

## Introduction

As of March 24, 2020, the worldwide tracking for Covid-19 infections reported 1,603,433 cases including 95,716 deaths^*1*^. Among the 18 countries that have reported more than 20,000 Covid-19 cases, the total number of cases per population ranged from 3,277 per million people in Spain to 57 per million cases in China and the total death rate among these countries ranged from 330 per million in Spain to 2 per million in China^*1*^. There are numerous potential factors to explain the wide variability in the number of infections and death among the countries including stringency of social isolation and contact tracing, the importation of the SARS-Cov-2, intensity of testing for the virus, as well as the preparedness and capacity of health care system to cope with the pandemic.

Differences among the host, particularly the sex of the patient and presence of selected co-morbid diseases and immunosuppressive state have been identified as risk factors from severe Covid-19^2,3^. Pre-existing medical conditions have been associated with increased prevalence of death including: cardiovascular disease (13.2%); diabetes (9.2%); chronic respiratory disease (8.0%); hypertension (8.4%) and recent diagnosis of cancer (7.6%) ^*4*^. Specifically, hypertensive patients had a hazard ratio of 1.70 for death ^*5*^ and 3.05 for in-hospital mortality ^*4*^. An increased association with the use of angiotensin-converting enzyme inhibitors (ACEIs) and angiotensin II receptor blockers (ARBs) has been reported for severe Covid-19 cases^*6*^. A biologically plausible link between SARS-CoV-2 infections and ACE inhibition has been proposed ^7^.

Angiotensin-converting enzyme 2 (ACE2) cleaves peptides within the renin-angiotensin system ^*8*^. ACE2 is mostly expressed in kidneys and the GI tract, and smaller amounts are found in type 2 pneumocytes in the lung and peripheral blood. The ACE2 receptor serves as the host cell entry for SARS-CoV and more recently it has been shown to be receptor for cellular entry for SARS-CoV-2^9,10^. As recently reported by *Hoffman et al* ^*11*^, the cellular entry of SARS-CoV-2 can be blocked by an inhibitor of the cellular serine protease TMPRSS2, which is employed by SARS-CoV-2 for S protein priming. Further attention has been drawn to the ACE2 receptor as antimalarials, which can interfere with ACE2 expression, have resulted in increased clearing of the SARS-CoV-2 clearance among Covid-19 patients ^*12*^.

To explore the variability in genetic polymorphisms and expression in human ACE2 (hACE2), we set out to determine if there were any differences between the Asian and Caucasian populations for ACE2 polymorphisms and compare the variability of hACE2 expression in peripheral blood among eight different populations.

### Comparison of genetic polymorphisms of hACE2 and its possible functional alteration between Asians and Caucasians

In order to investigate whether differences in genetic variations exist between Caucasians and Asians and if these variants can influence the efficiency of cell entry of SARS-CoV-2, we retrieved the variants in the hACE2 from gnomAD v2.1 exomes^13^. We used 56,885 Non-Finnish European (NFE) and 9,197 East Asians (EAS, including 1,909 Koreans) for our analysis. There were 5,693 mutations of which 177 were missense mutations. In comparing the allele frequency of Caucasian and Asian cohorts, Fisher’s exact t tests were performed and adjusted for multiple testing. Four genetic variants reached statistical significance (adjusted p-value less than 0.05), although their effect size was modest (Details in Table1). One variant was located at the a1 helix of ACE2 gene (K26R), two in the middle of SARS-CoV-2 (N638S and I468V) and one in the tail (N720D). The K26R variant was of immediate interest, since the spike glycoprotein (S protein) of virus contains the receptor binding domain (RBD), which directly binds to the peptidase domain (PD) of hACE2. The PD of hACE2 mainly engages the a1 helix in the recognition of the RBD^14^.

**Table 1.**
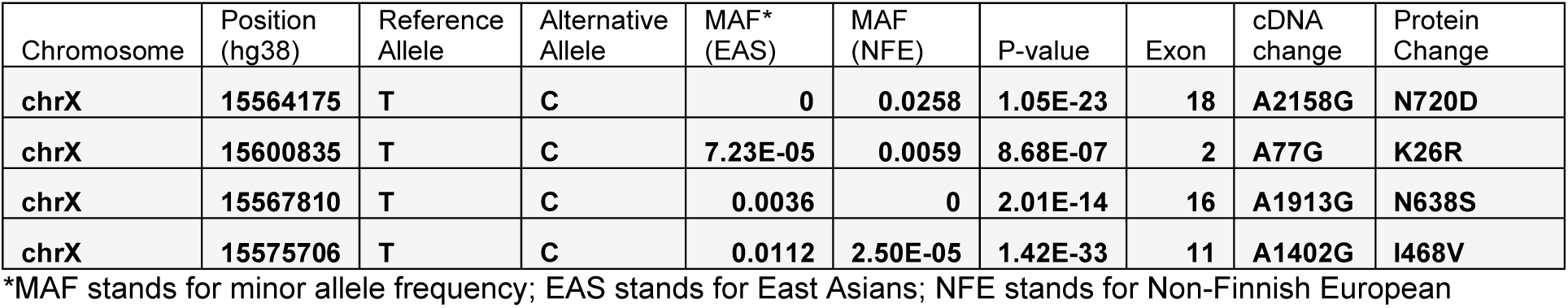
Four missense mutations in hACE2 show significant allele frequency difference between EAS and NFE

We also showed these four mutations in the hACE2 X-ray structure as Figure1.

**Figure 1.**
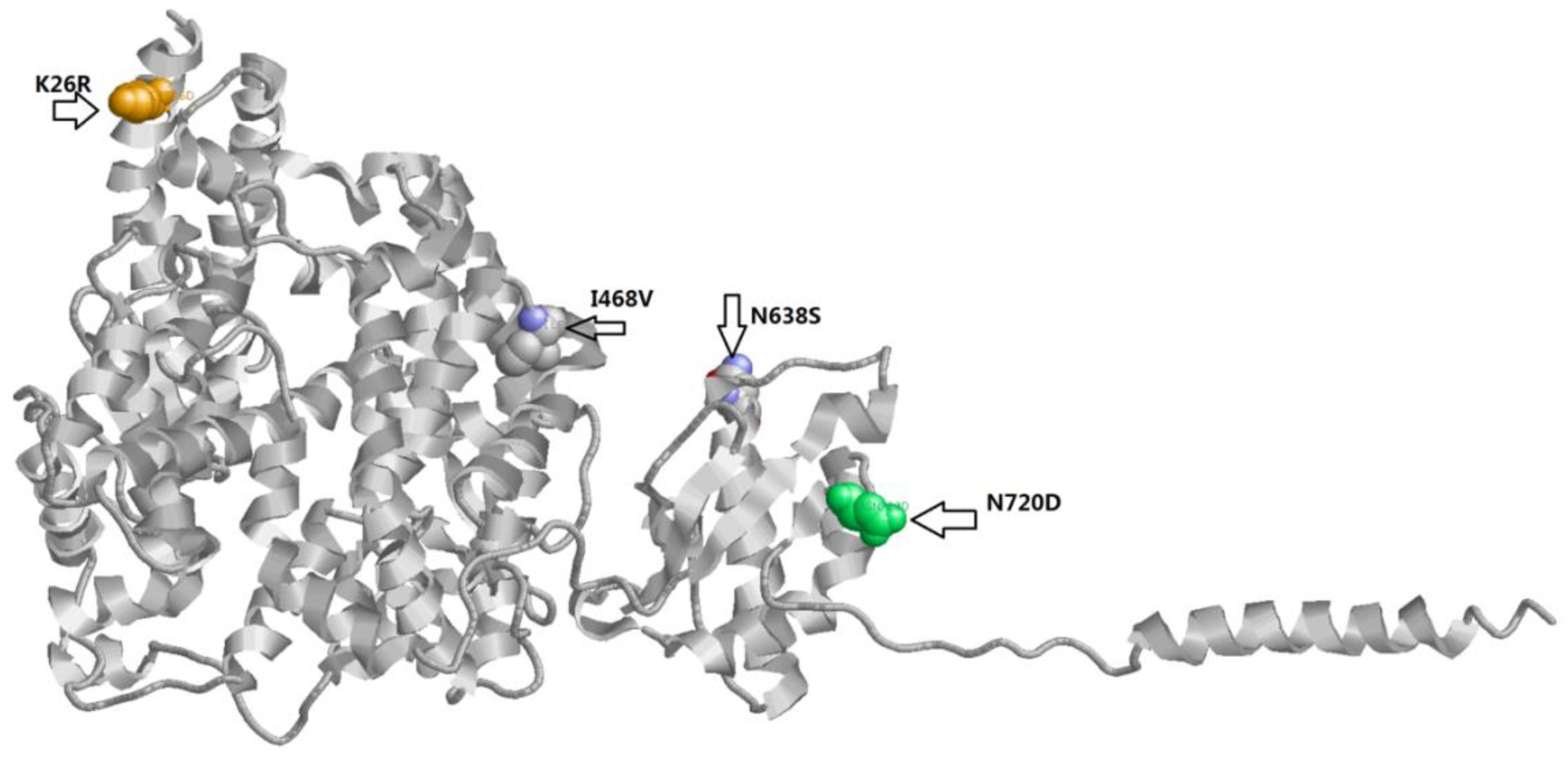
Human ACE2 structure and four missense mutations that differ between Caucasians and Asians.

Molecular dynamics (MD) simulation which is a canonical method for investigating the influences of mutant on protein structure, was then applied. As mutations N638S and N720 are far away from the RBD, as a result we mainly focused on the K26R as MD simulations demonstrated that this variant mutated more frequently in Non-Finnish European while I468V mutated frequently more in East Asians. The initial wild type structure of the human ACE2 (hACE2) was selected from the X-ray crystal complex structure between hACE2 with SARS-CoV-2 spike (S) protein from Protein Data Bank^15^ (PDB ID: 6LZG). The initial structures of mutants were constructed using FoldX (http://foldxsuite.crg.eu/command/PositionScan). The structural minimizations and molecular dynamics simulation was performed using the GROMACS 2018.4^16-18^ software package. OPLS-AA force field was adopted for the protein complex and the SPCE water model for solvate the system. The periodic boundary conditions (PBC) in a cubic box was applied, and the distance to the edge of the box was set as 1.0 nm. After the protein complex system was solvated, the appropriate numbers of sodium and chloride ions were added to neutralize the system. Equilibration simulations were done under the NVT ensemble for 2 ns and then the NPT ensemble for 1 ns after an extensive energy minimization by using the steepest descent algorithm. The simulation time step was 2 fs. The temperature was maintained at 300K using the V-rescale algorithm with a tau-t of 0.1 ps. The pressure was maintained at 1 atm by the semi-isotropic Parrinello−Rahman method with a tau-p of 2 ps and a compressibility of 4.5×10-5 bar-1. The Particle Mesh Ewald (PME) method was used for calculation the long-range electrostatic interactions.

The LINCS algorithm was used for constraining all covalent bonds. Finally, a 100ns long MD simulation was carried out for each system. The stability of the simulation was checked by computing root mean square deviations (RMSDs). RMSDs of Cα were calculated for the mutant K26R/I468V and compared to the wild-type shown in Figure 2. RMSDs for the complexes with K26R changed little during the 100ns MD simulations. Mutant I468V showed RMSD increase during time 20-30 ns, but no obvious RMSD change comparing with wild type after 40ns. The RMSD values for the K26R mutant are slightly higher and structure change compared with wild type.

**Figure 2.**
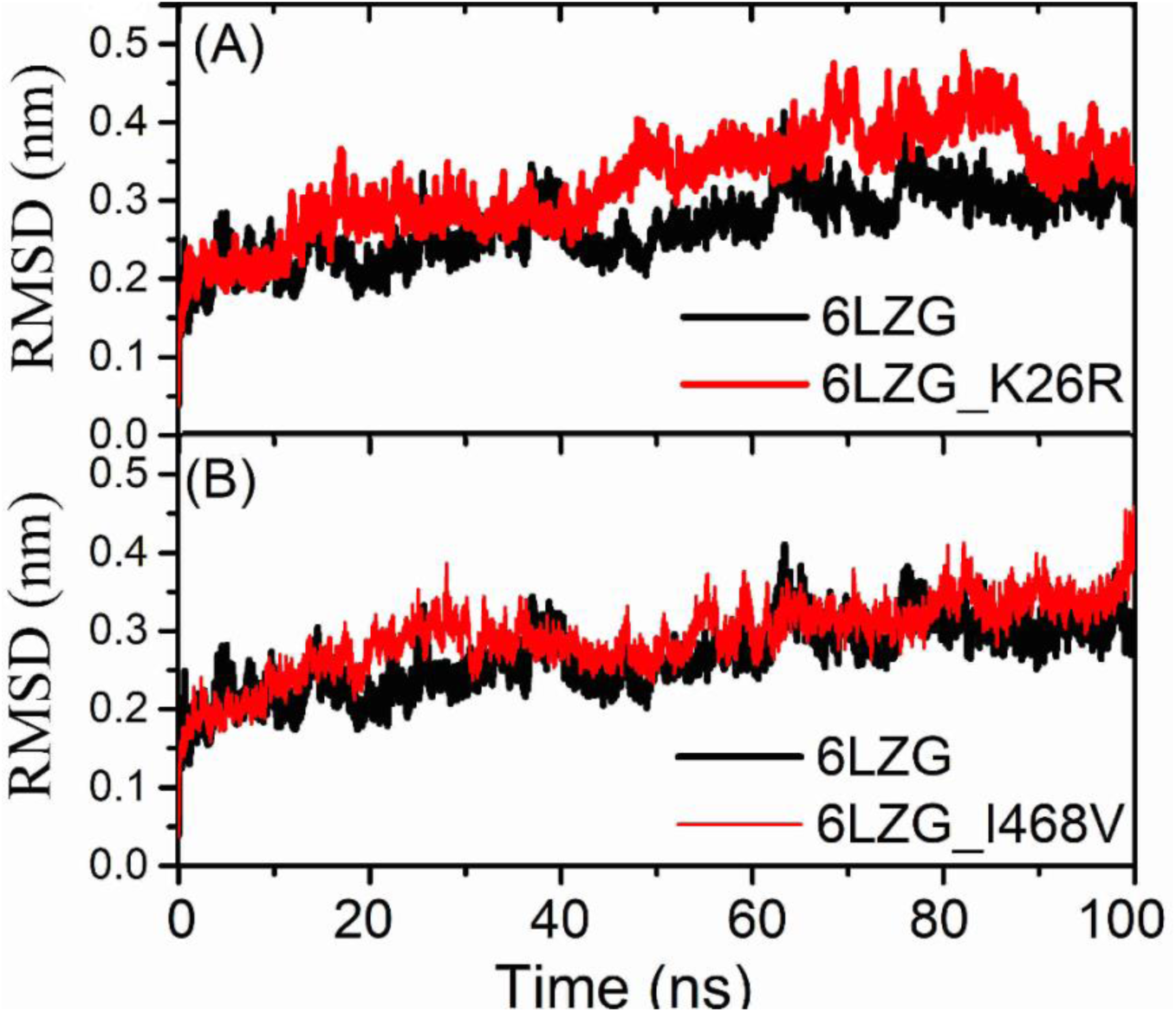
RMSDs of hACE2 wild type (black line) and mutated hACE2 protein(red line) during Molecular dynamics simulation.

The effect of missense on protein-ligand complex can be assessed by various experimental technologies^19^, but are very time-consuming. The *in silico* binding free energies calculation can be performed to determine the effects of mutations. Here, the binding free energies between S protein and hACE2 (wild type or mutant) were calculated using the g_mmpbsa (Table 2). These two mutations K26R/I468V were both predicted to slightly increase the binding free energy and may slightly decrease the binding affinity. While, K26R mutated more frequently in Caucasian, and I468V mutated more frequently among Asians.

**Table 2.**
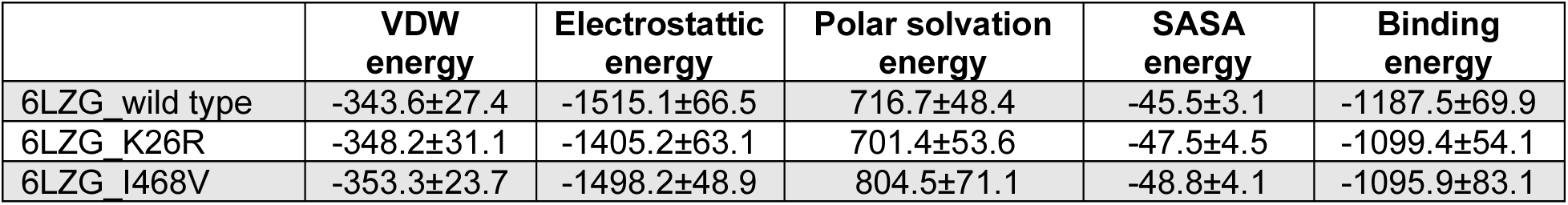
Binding free energy (in units of kJ/mol) for S protein and hACE2 (wild type and mutated) complex

### Variability in gene expression of hACE2 gene among various ethnic populations

The second goal was to assess the expression variability of hACE2 in eight global populations from the HapMap3 project^20,21^. The following eight populations were compared: Utah residents with Northern and Western European ancestry from the CEPH collection (CEU); Han Chinese in Beijing China (CHB); Gujarati Indians in Houston, Texas(GIH), Japanese in Tokyo, Japan (JPT); Luhya in Webuye, Kenya (LWK); Mexican ancestry in Los Angeles (MEX), California, Maasai in Kinyawa, Kenya (MKK); and Yoruba in Ibadan, Nigeria (YRI). Raw expression data from Sentrix Human-6 Expression BeadChip version 2 were extracted, background-corrected, log2 scaled and quantile normalized. Expression data from the populations with admixture (GIH, LWK, MEX, and MKK) were also normalized for population admixture using EIGENSTRAT^22^. The normalized expression data were used for the assessing the expression level for different sets of analysis. The hACE2 expression levels of these 8 populations were showed in Figure-3.

**Figure 3.**
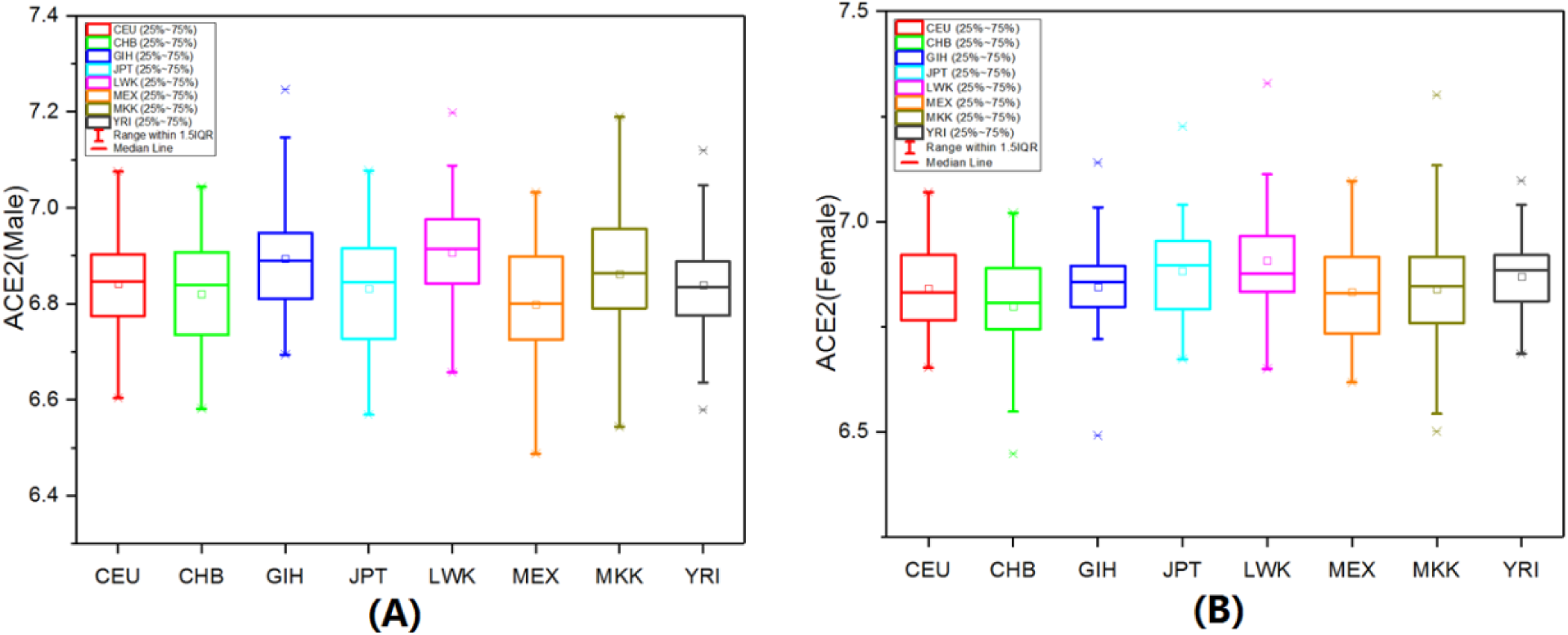
hACE2 expression among 8 Hapmap3 populations. A) for Males and B) for Females.

The hACE2 expression level of CHB population (42 females and 38 males) was then compared to other cohorts using ANOVA analysis, after stratification for sex(Table-3). Marginal differences were noted among the populations. There was a statistically higher hACE2 expression in men for LWK and GIH and JPT populations, LWK and YRI populations for females. However the effect sizes were small and its clinical relevance is not clear.

**Table 3.**
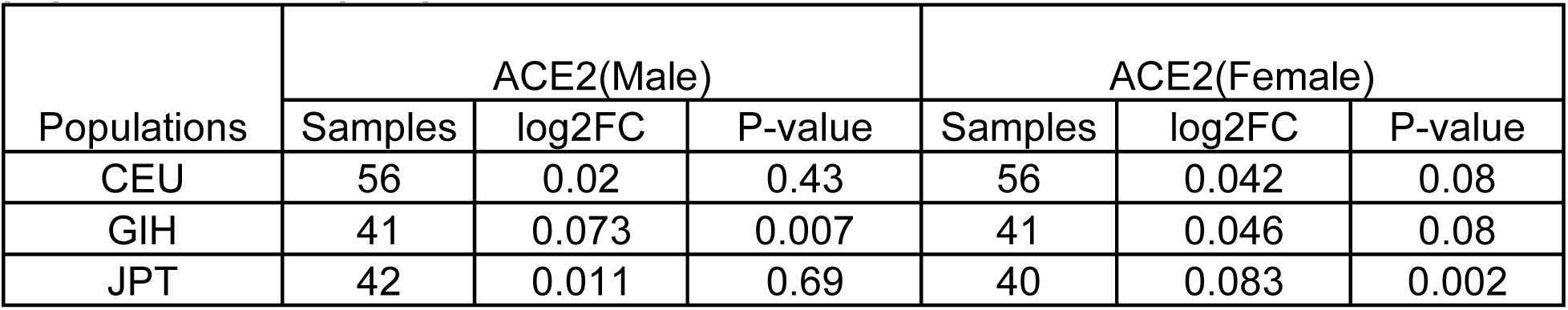
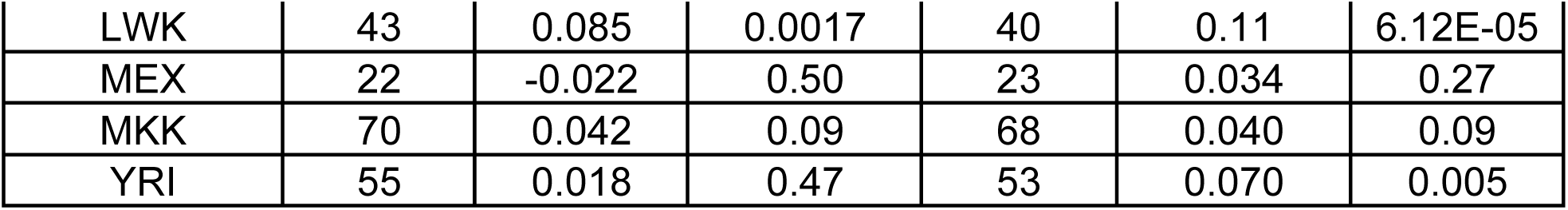
Comparison of ACE2 expression between Asians (CHB) and other 7 populations in Hapmap3

| Populations | ACE2(Male) |  |  | ACE2(Female) |  |  |
| --- | --- | --- | --- | --- | --- | --- |
|  | Samples | log2FC | P-value | Samples | log2FC | P-value |
| CEU | 56 | 0.02 | 0.43 | 56 | 0.042 | 0.08 |
| GIH | 41 | 0.073 | 0.007 | 41 | 0.046 | 0.08 |
| JPT | 42 | 0.011 | 0.69 | 40 | 0.083 | 0.002 |
| LWK | 43 | 0.085 | 0.0017 | 40 | 0.11 | 6.12E-05 |
| MEX | 22 | -0.022 | 0.50 | 23 | 0.034 | 0.27 |
| MKK | 70 | 0.042 | 0.09 | 68 | 0.040 | 0.09 |
| YRI | 55 | 0.018 | 0.47 | 53 | 0.070 | 0.005 |

## Discussion

This study that examined the genetic polymorphisms in the hACE2 gene between Caucasians and Asians. We noted that there were four missense mutations, of which K26R and I468V were of most interest based on their location. The propensity for these variants to mutate were different among these two polymorphisms. Among Caucasians, K26R mutated more, while among Asians I468V mutated more frequently. In silico studies have noted that these two mutations may affecting the binding characteristics between S protein and hACE2, however in the absence of further functional studies, the significance of these alternations is not clear.

We also found that there was some variability of genetic expression of hACE2 among various populations, but the magnitude of the differences was small and so it is unclear if this has any impact based on this subtle differences in susceptibility or severity of Covid-19 among various ethnicities. We also did not notice a differential expression of hACE2 between the sexes, even though it is widely reported that males have a worse prognosis than females for complications of Covid-19 ^*4*^. Our findings are generally consistent with a recently published study by *Chen et al* ^*23*^, that reported the ACE2 expression in Asians was similar to that of other races. Our expression study was conducted in peripheral blood and the results for site specific expression (alveolar, GI or renal tissue) may yield different results.

In conclusion, our results do reveal some differences in genetic polymorphisms between Asians and Caucasians which may potentially alter the binding of the virus to the hACE2 receptor and some subtle variation in genetic expression of hACE2 among different populations. However our findings need to replicated and further in vitro studies should performed, before the significance of these findings can be determined. At present the reasons behind the variability in Covid-19 presentation among the various countries needs to be further explored, as we were unable to provide convincing evidence to suggest there are differences in allele frequency or expression of hACE2 among Caucasians and Asians.

## Compliance with Ethical Standards

Conflict of Interest: None of the authors have no conflicts of interest to disclose regarding this manuscript.

## Human and Animal Rights and Informed Consent

Data used for this study was obtained from public databases.

